# Integrative taxonomy confirms the species status of the Himalayan langurs, *Semnopithecus schistaceus* Hodgson 1840

**DOI:** 10.1101/602243

**Authors:** Kunal Arekar, S. Sathyakumar, K. Praveen Karanth

## Abstract

Taxonomy is replete with groups where the species identity and classification remain unresolved. One such group is the widely distributed Hanuman langur (Colobinae: *Semnopithecus*). For most part of the last century, Hanuman langur was considered to be a single species with multiple subspecies. Nevertheless, recent studies using an integrative taxonomy approach suggested that this taxon is a complex, with at least three species. However, these studies did not include the Himalayan population of the Hanuman langur whose taxonomic status remains unresolved. The Himalayan population of Hanuman langurs have been classified as a distinct species with multiple subspecies or have been subsumed into other species. These classification schemes are wholly based on morphological characters and which are sometimes insufficient to delimit different species. Here, we have integrated data from multiple sources viz. morphology, DNA, and ecology to resolve the taxonomy of the Himalayan langur and to understand its distribution limit. Our results with three lines of evidence corresponding to three different species concepts show that Himalayan langur is a distinct species from *S. entellus* of the plains. Additionally, these results did not show any support for splitting of the Himalayan langur into multiple subspecies. Our study supports the classification proposed by Hill (1939) and we recommend *Semnopithecus schistaceus*, Hodgson 1840 as species name for the Himalayan langur and subsume all the known subspecies into it.

## Introduction

Species is the fundamental unit of study in many fields of biology such as systematics, ecology, evolution, behaviour and many more (de Queiroz, 2007). Thus, the first step in these studies is accurate and unambiguous identification of species for which it is imperative to have a well resolved taxonomy of the group of interest. Nevertheless, taxonomy is replete with “problematic groups” wherein species identity and classification remain unresolved. One such group are the colobine monkeys broadly referred to as Hanuman langur (Genus: *Semnopithecus*, Subfamily: Colobinae). Hanuman langur is a widely distributed primate in the Indian subcontinent (Bishop, 1977; Newton, 1988) which exhibits extensive morphological variation across its range. A multitude of classification schemes have been proposed mostly during early-mid 20^th^ century to resolve the taxonomic status of Hanuman langurs (Brandon-jones, 2004; Ellerman, J. R. and Morrison-Scott, 1966; Groves & C, 2001; Hill, 1939; Napier & Napier, 1967; Pocock, 1928, 1939; Roonwal, 1984; Roonwal & Mohnot, 1977).

Hanuman langur is broadly divided into two categories; the northern type (NT) which is characterised by forward looping tail towards the head and southern type (ST) with the tail looping backwards away from the head (Roonwal, 1979, 1984). The Tapti-Godavari rivers in central India form the borderline with NT distributed to the north and ST distributed to the south of these rivers (Roonwal, 1979, 1984). Recent studies support the splitting of ST Hanuman langur into two species, namely *S. priam* and *S. hypoleucos*, based on an integrative approach wherein multiple lines of evidence from molecular, morphological and ecological data were used (Ashalakshmi, Nag, & Karanth, 2014; Nag, Karanth, & Vasudeva, 2014; Nag, Pramod, & Karanth, 2011). Similarly, genetic and morphological data suggest that the plains population of NT Hanuman langur is a separate species, *S. entellus* (Karanth, Singh, & Stewart, 2010). However, taxonomy of the Himalayan population (hereafter Himalayan langur) is still unresolved. In order to understand the exact number of species of Hanuman langur in the Indian subcontinent, it is important to resolve the taxonomy of all the populations of Hanuman langur.

Himalayan langur is the northernmost population of Hanuman langur (Sugiyama, 1976) distributed in the Himalayan region of India, Nepal and parts of Pakistan and Bhutan (Blanford, 1888; Pocock, 1939). Altitudinally, they are distributed from the Himalayan foothills up to 4270m asl (above sea level) (Bishop, 1977) and is one of the few colobine monkey living in temperate climate (Nijman, 2013), while the rest are distributed predominantly in tropical regions (Bishop, 1979). Himalayan langurs are classified in the NT category based on the tail loop character (Roonwal, 1979, 1984). Behavioural characters suggests that the Himalayan langurs might be distinct from *S. entellus* of the northern plains. (Bishop, 1975; Bishop, 1979; Curtin, 1982; Sugiyama, 1976), Morphologically, the Himalayan langur can be distinguished from the plains’ population *(S. entellus)* by the tail carriage pattern (discussed in methods section) and by their pelage – the langurs from the Himalayas have a bushy white head which is very distinct from the darker grey-brown body (Bishop, 1979).

A variety of classification schemes have been proposed to resolve the taxonomy of the Himalayan langurs (Table 1). One of the earliest comprehensive classification of Indian colobines was by Pocock (1928). He assigned the Himalayan langurs to five subspecies *schistaceus, ajax, achilles, lanius* and *hector* under the species *Pithecus entellus*. Later Pocock (1939) renamed *Pithecus* as *Semnopithecus* with three subspecies under it; *schistaceus, ajax* and *achilles*. The subspecies *lanius* and *hector* from Pocock’s (1928) earlier classification were not included here. Hill (1939) considered the Himalayan langur to be a single species *S. schistaceus* with five subspecies *hector*, *lanius*, *achilles*, *schistaceus* and *ajax*.

**Table 1:** Different Taxonomic Schemes (TS) for Himalayan langur proposed by various

| TS 1 |  |  |  |  | TS 2 | TS 3 |
| --- | --- | --- | --- | --- | --- | --- |
| Pocock (1928) | Pocock (1939) | Roonwal and Mohnot (1977) | Roonwal (1984)* | Brandon-Jones (2004) | Hill (1939) | Groves (2001) |
| *Pi. e. schistaceus* | *S. e. schistaceus* | *P. e. schistaceus* | *P. e. schistaceus* | *S. e. schistaceus* | *S. schistaceus* | *S. schistaceus* |
| *Pi. e. ajax* | *S. e. ajax* | *P. e. ajax* | *P. e. ajax* | *S. e. ajax* | *S. s. ajax* | *S. ajax* |
| *Pi. e. achilles* | *S. e. achilles* | *P. e. achilles* | *P. e. achilles* | — | *S. s. achilles* | — |
| *Pi. e. lanius* | — | *P. e. lania* | *P. e. lania* | — | *S. s. lanius* | — |
| *Pi. e. hector* | — | — | — | *S. e. hector* | *S. s. hector* | *S. hector* |

Subsequent classification schemes (Ellerman and Morrison-Scott, 1966; Napier & Napier, 1967; Roonwal, 1984; Roonwal & Mohnot, 1977) synonymised *Semnopithecus* with *Presbytis* and subsumed all the Himalayan species into a single species *P. entellus* along with the subspecies *entellus* from the northern plains. The subspecies *lanius* (Hill, 1939) was changed to *lania* and the subspecies *hector* (Hill, 1939) was removed.

Later, Groves (2001) reverted back to using *Semnopithecus* for Hanuman langurs and recognized three species of Himalayan langur *S. schistaceus, S. ajax* and *S. hector*. He elevated the three subspecies from previous classification schemes to species level. Lastly, Brandon-Jones (2004) included all the Himalayan species as subspecies of *S. entellus*, except for *achilles* and *lanius* (Hill, 1939) which he did not include in the classification.

Thus, the Himalayan langur has had a convoluted taxonomic history which fall into three broad groups of taxonomic schemes (TS). TS1 – Various populations of Himalayan langurs are placed in multiple subspecies under *Pi. entellus, S. entellus* or *P. entellus* (Pocock, 1928, 1939; Ellerman and Morrison-Scott, 1966; Napier and Napier, 1967; Roonwal and Mohnot, 1977; Roonwal, 1984; Brandon-Jones et al., 2004). TS2 – Himalayan langurs is considered a distinct species with multiple subspecies (Hill, 1939). TS3 – Himalayan langurs is split into multiple species (Groves, 2001).

With the advent of molecular techniques many recent studies have used genetic data to resolve taxonomic ambiguities in primates of the Indian subcontinent (Arekar, Parigi, & Karanth, 2019; Ashalakshmi et al., 2014; Chakraborty, Ramakrishnan, Panor, Mishra, & Sinha, 2007; Karanth, Singh, Collura, & Stewart, 2008; Karanth et al., 2010; Osterholz, Walter, & Roos, 2008; Wangchuk, Inouye, & Hare, 2008). However, the use of molecular data does not guarantee a robust description and identification (Will, Mishler, & Wheeler, 2005). Molecular data is often considered as another singular data type, such as the morphological data, which can be used as a line of evidence to characterise and describe species. In order to achieve a robust delineation of species, we need to integrate methods from different discipline (Dayrat, 2005).

Given this background, we have used data from multiple sources viz. morphology, DNA and ecology to address the following questions 1) Is Himalayan langur a distinct species from *S. entellus*? 2) Does Himalayan langur occupy different niche than *S*. entellus? 3) Does Himalayan langur population consist of multiple species/subspecies? And 4) What is the potential distribution range of the Himalayan langurs.

## Materials and Methods

### Molecular data

#### Data collection

We conducted field work in four Himalayan states of India – Jammu and Kashmir (J&K), Himachal Pradesh (HP), Uttarakhand and Sikkim. These states were chosen for field work based on distribution records from past studies (Pocock, 1939; Hill, 1939; Sugiyama, 1975; Bishop, 1979: Choudhury, 2001) as well as from IUCN website (www.iucnredlist.org). Furthermore, we included our field data and published sequences from Nepal.

Molecular data was based on 175 faecal samples collected from 46 locations (Table S1) from across the distribution range of Himalayan langur in India and which included multiple samples collected from each location. Additionally, five faecal samples of *S. entellus* were collected from one location in the northern plains (22.88220N, 88.39970E). Fresh faecal samples were collected by following the troops in the morning and the evening hours. Samples were collected by two different methods – First, as described in Kawamoto et al. (2013), a sterile cotton swab was rolled multiple times over the surface of the faeces and thoroughly rinsed in the lysis buffer (White & Densmore, 1992). Second method involved collection of the whole faeces which was then stored in absolute alcohol. These samples were stored at −20 °C in the laboratory until DNA extraction. Samples stored in lysis buffer were first treated with starch to remove potential PCR inhibitors like bilirubin and bile salts (Kawamoto et al., 2013; Zhang, Li, Ma, & Wei, 2006) and then DNA was extracted by using Wizard^®^ SV Gel and PCR Clean-Up System, Promega, Madison, WI, USA and stored in pure nuclease free water at −20 °C until further use. DNA from whole faecal samples was extracted using the commercially available QIAamp DNA stool mini kit (QIAGEN Inc.), following the manufacturer’s protocol with slight modifications as mentioned in Mondol et al. (2009), however, we did not add the carrier RNA (Poly A) (Kishore, Reef Hardy, Anderson, Sanchez, & Buoncristiani, 2006;). Each extraction had a negative control to monitor contamination. The quantity of extracted DNA was measured using a NanoDrop 2000 Spectrophotometer (Thermo Fisher Scientific Inc).

#### PCR amplification and sequencing

A 700 bp region of mitochondrial cytochrome *b* (Cyt-*b*) gene was PCR amplified using the primer pair Cytb_278F (5’ – GCCTATTTCTACACGTAGGCCG – 3’) and Cytb_1052R (5’ – CCAATTGCAATGAAGGGTTGGT – 3’). A 25 μl of reaction was set with standard 1X reaction buffer premixed with 1.5 mM MgCl_2_ (New England BioLabs^®^ Inc.), 0.25mM of dNTPs (Bangalore Genei, Bangalore), 0.3 μM of each primer (Sigma-Aldrich Inc.), 1.5U Taq polymerase (New England BioLabs^®^ Inc.) and 2 μl of template DNA (DNA concentration between samples varied from 20 ng/μl to 80 ng/μl). We also added 2 μl of BSA (Bovine Serum Albumin, Fisher Scientific) to augment the PCR reaction. Further, the template DNA was diluted to 1:5 (DNA extract: water) ratio and used for the reaction in order to reduce the amount of PCR inhibitors. The PCR cycling conditions were carried out with initial denaturation at 94 °C for 5 mins followed by 50 cycles with denaturation at 94 °C for 40 sec, annealing at 57.5 °C for 30 sec and elongation at 72 °C for 30 sec and final extension at 72°C for 10 mins. The PCR products were outsourced for purification and sequencing to Medauxin, Bangalore.

#### Phylogenetic analysis

The sequence files obtained were viewed and edited manually in ChromasLite v2.01 (Technelysium Pty Ltd). Sequences Himalayan langur as well as *S. entellus* were also downloaded from a previous molecular study (Supplementary material, Table S5) (Ashalakshmi et al., 2014; Khanal, Chalise, Wan, & Jiang, 2018). The sequences were aligned using MUSCLE algorithm (Edgar, 2004) incorporated in MEGA v7 (Kumar, Stecher, & Tamura, 2016).

We used jModelTest 2.1.3 (Darriba, Taboada, Doallo, & Posada, 2012) to pick the best model of sequence evolution. Phylogenetic reconstruction was performed using Maximum Likelihood (ML) and Bayesian methods. ML analysis was performed in RAxML7.4.2 incorporated in raxmlGUI v1.3 (Stamatakis, 2006). We used GTR+G model in RAxML as there is no provision to use other models. 1000 replicates were performed to assess support for different nodes. We used MrBayes 3.2.2 (Ronquist et al., 2012) to perform the Bayesian analysis with HKY+G nucleotide substitution model. Two parallel runs with four chains each were run for 10 million generations with sampling frequency every 1000 generations. Convergence between the two runs was determined based on standard deviation of split frequencies. The program Tracer v1.6 (Rambaut, A., Suchard, M.A., Xie, D., Drummond, 2013) was used to determine stationarity, an effective sample size (ESS) value of >200 for each parameter was used as a cut-off for run length. The first 25% of trees were discarded as burn-in.

#### Hypothesis testing

We compared the Bayesian tree (Fig. 1), with an *a priori* hypothesis where the phylogeny was constrained to be consistent with Groves’ (2001) three species of Himalayan langurs; *S. ajax, S. hector* and *S. schistaceus*. RAxML7.4.2 incorporated in raxmlGUI v1.3 (Stamatakis, 2006) was used to generate a constraint tree. The likelihood of the constraint tree was then compared with the best tree using the SH test (Shimodaira & Hasegawa, 1999) and AU test (Shimodaira, 2002) in PAUP* Version 4.0a (build 164) (Swofford, 2001). For both the SH and AU test, we performed 10000 bootstrap replicates and the nonparametric bootstrap with reestimated log likelihoods (RELL) approximation (Kishino, Miyata, & Hasegawa, 1990) was used for resampling the loglikelihoods.

### Morphological data

#### Data collection

Morphological data was collected by direct observations during field survey as well as from photographs. Based on past studies (Nag et al., 2011) and our field observations, six color-independent morphological characters were used to differentiate between *S. entellus* and the Himalayan langur. These included the four characters described by Nag et al. (2011) and two characters unique to Himalayan langurs. Characters specific to Himalayan langur included tail carriage (Roonwal, 1979, 1984) and demarcation between the head and the body (HBC) (Bishop, 1979; Groves, 2001; Hill, 1939; Pocock, 1939). Within the NT langurs, two forms of tail carriage are observed, in *S. entellus* from the northern plains the tail loops over the back and the tip of the tail hangs perpendicular to the ground, here onwards TC3 (Fig. 1b), whereas, in the Himalayan populations the tail loops well behind the back and the tip ends above the base of the tail, here onwards TC4 (Fig. 1a). We coded them TC3 and TC4 to be consistent with Nag et al.’s (2011) coding system. The tail carriage pattern was recorded when the individual was walking on a flat surface and not while climbing up or down the hill and nor while it was running or standing (as per Roonwal, 1984). The Himalayan langur has a distinct demarcation between the head and the body, the head is bushy and white coloured distinct from the grey-brown body (Fig. 2a) (Bishop, 1979; Groves, 2001; Hill, 1939; Pocock, 1939). On the other hand, the langurs from the plains (*S. entellus*) have a uniform colour without any distinction between the head and the body (Fig. 2b). We call this character Head-Body contrast (HBC) character. Apart from these two characters we also used the four characters described in Nag et al. (2011), these characters are not seen in Himalayan langurs and therefore were coded ‘0’(absent). We recorded these characters by direct observations using 10 × 50 binoculars (Olympus) and through photographs taken from a digital camera (Canon PowerShot SX20 IS). These characters were scored for multiple adult individuals per location.

**Fig. 1:**
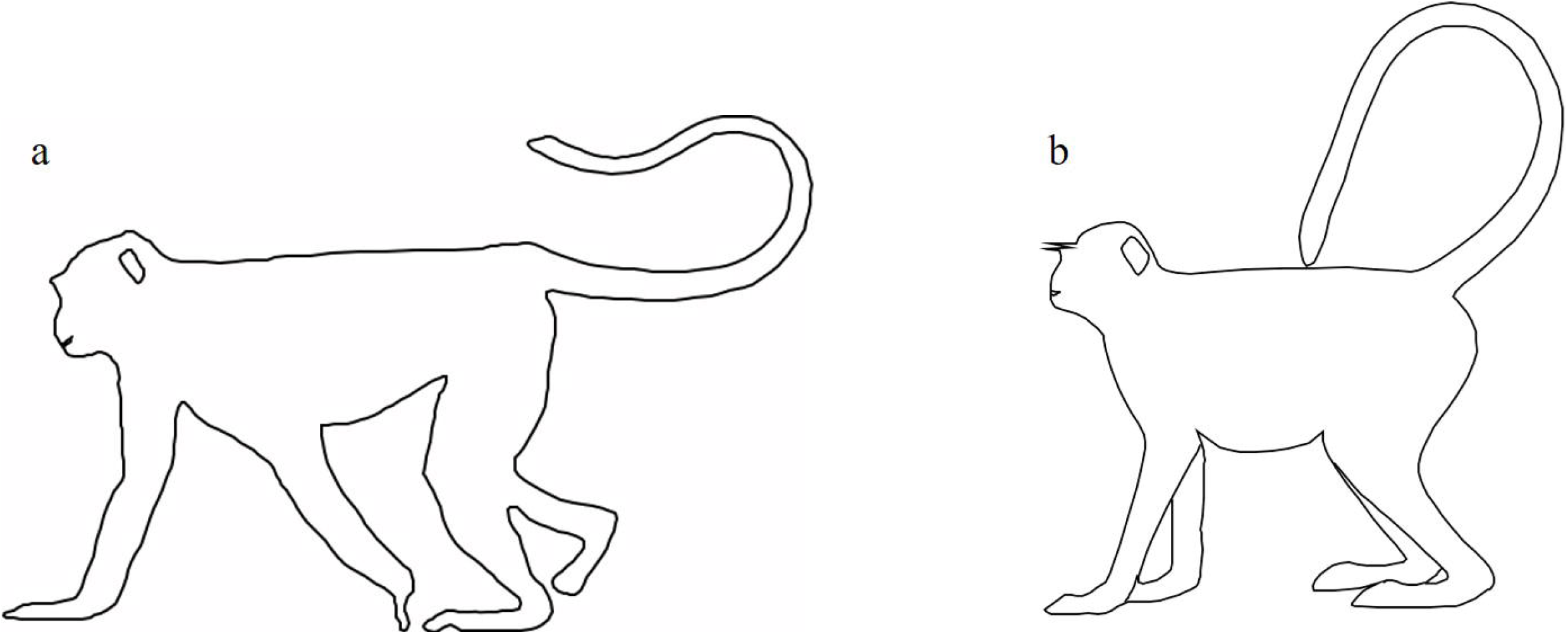
Tail carriage pattern in the Himalayan langur (a) and *Semnopithecus entellus* (b).

#### Analysis

We typed 65 adult individuals from 29 locations (Supplementary material, Table S6). We also included one of the southern species from this complex, *S. hypoleucos* (St1 and St2 morphotypes in Nag et al., 2011) in the analysis. The characters were coded as described in Table 2. All the characters were coded in a way that they are consistent with Nag et al.’s (2011) coding system. The codes for *S. hypoleucos* were similar to ones used in Nag et al. (2011).

**Table 2:** Coding system used for the 6 morphological characters used in this study. The coding system is designed to be consistent with Nag et al. (2011), unless mentioned otherwise.

|  | Characters |  |  |  |  |  |
| --- | --- | --- | --- | --- | --- | --- |
|  | Crest | Streak | EOB | Tail loop<br>(TL) | Tail carriage<br>(TC) | HBC |
| Himalayan<br>langur | 0 | 0 | 0 | 1 | 3 | 1 |
| *Semnopithecus*<br>*entellus* * | 0 | 0 | 3 | 1 | 4<sup>#</sup> | 0 |
| *Semnopithecus*<br>*hypoleucos* | 0 | 1 | 3/4<sup>†</sup> | 2 | 1 | 0 |
| *Semnopithecus*<br>*priam* ** | 1 | 0 | 1 | 2 | 2 | 0 |

\**S. entellus* = Nt from Nag et al. (2011).
<sup>#</sup>Nag et al. (2011) coded Nt as 0, here we have coded it 4.
<sup>†</sup>*S. hypoleucos* contains two morphotypes (St1/St2; Nag et al., 2011).
\*\* We did not use this species in the analysis.

We prepared a character matrix with the terminal taxa in the rows and the six morphological characters in the columns. Using this matrix, first the mean character difference was calculated between the individuals in PAUP* Version 4.0a (build 164) (Swofford, 2001). Then a Neighbour Joining (NJ) tree was built using these distances, with the default settings. Mid-point rooting was used to root the tree.

### Ecological data

#### Data collection

For ecological data, we obtained 192 occurrence records of the Himalayan langurs (Available with authors on request). Out of these, 79 records were from the field surveys conducted for this study, 58 occurrence records were obtained from previous studies (Khanal et al., 2018; R.A. Minhas et al., 2018; Minhas et al., 2012) and 55 occurrence records were downloaded from GBIF (Global Biodiversity Information Facility) database (www.gbif.org). For *S. entellus*, 69 occurrence records were used, out of these, 21 records were obtained from Nag et al. (2014) and 48 records were downloaded from GBIF database (www.gbif.org). For the occurrence records downloaded from GBIF database, we plotted these occurrence records on the map and included only those records which were falling within the known distribution zones of the respective taxa. Further, we used 22 environmental layers, 19 were bioclimatic layers downloaded from www.worldclim.org, one altitude layer (USGS website), two more layers – slope and aspect were derived from the altitude layer in ArcGIS 10.2.1 (Table S2). All the layers were of 30 arcsec resolution, projected in WGS84 projection. These bioclimatic variables were clipped to the region from 68 °E to 97.4 °E and from 6.7 °N to 37 °N using ArcGIS 10.2.1. These clipped layers were then exported to ASCII format using QGIS 2.18.12. The 22 environmental layers were tested for multicollinearity by calculating Pearson’s correlation coefficient (r). The layers with r ≤ 0.75 were selected for further analysis (Supplementary material, Table S3).

#### Ecological Niche Modelling analysis

##### Model selection

We first performed model selection using the maximum entropy algorithm available in Maxent v3.4.1 (Phillips, Anderson, & Schapire, 2006). We tested 48 models, each for Himalayan langur and *S. entellus*, by employing different combinations of features and regularisation multiplier (RM) (Supplementary material, Table S4) in MaxEnt v3.4.1 (Phillips et al., 2006).

##### MaxEnt algorithm

Separate Maxent analyses for the Himalayan langur and for *S. entellus* were implemented in MaxEnt v3.4.1. Maxent was run with the following modifications. Random test percentage was set to 30%, maximum number of background points was set to 10000 and the replicates were set to 10 with replicated run type changed to Subsample. 5000 iterations were performed with the convergence threshold set to 1 x 10^−5^. Jackknife test was used to estimate the contribution of each environmental variable. The feature type and RM value were based on the best model selected (Table S4). To overcome the sampling bias, a bias file was created in ArcGIS 10.2.1 by applying Gaussian kernel density function to 10000 background points (Elith, Kearney, & Phillips, 2010). The output format was chosen as Cloglog (Phillips, Anderson, Dudík, Schapire, & Blair, 2017). AUC values were examined to check for the predictive ability of the model.

Schoener’s D (Schoener, 1968) was calculated as a measure of niche overlap between the two distribution models (Warren & Seifert, 2011) implemented in ENMTools (Warren, Glor, & Turelli, 2010)

## Results

### Phylogenetic analysis

Out of the total 182 samples subjected to DNA extraction we obtained sequences from 76 samples. We also performed DNA extraction and sequencing for one *S. hypoleucos* sample, it was used as an outgroup in this study. These 76 sequences contained multiple identical sequences, so for building the phylogenetic tree, we trimmed the dataset to one unique sequence per location. Therefore, our final dataset contained 746 bp of Cyt-*b* sequence from 57 samples (Table S5), which included sequences from previous studies.

The Bayesian and ML analysis recovered two major clades, the *S. entellus* clade – containing the sequences from the northern plains; and the Himalayan clade – containing sequences from the Himalayan region (Fig 3). Within the Himalayan clade was a well-supported subclade consisting of haplotypes from the western Himalayas, whereas sequences from eastern Himalayas did not form a separate cluster. Both, the Bayesian and ML trees, showed similar topology where in all the major clades were retrieved. Furthermore, in the Bayesian tree (Fig. 1), two samples i.e.128_Nepal and 129_Nepal were placed within the clade containing samples from the Indian Himalayan Region (IHR), whereas in the ML tree (Supplementary material, Fig. S1), these two samples were sister to the above mentioned clade.

**Fig. 2:**
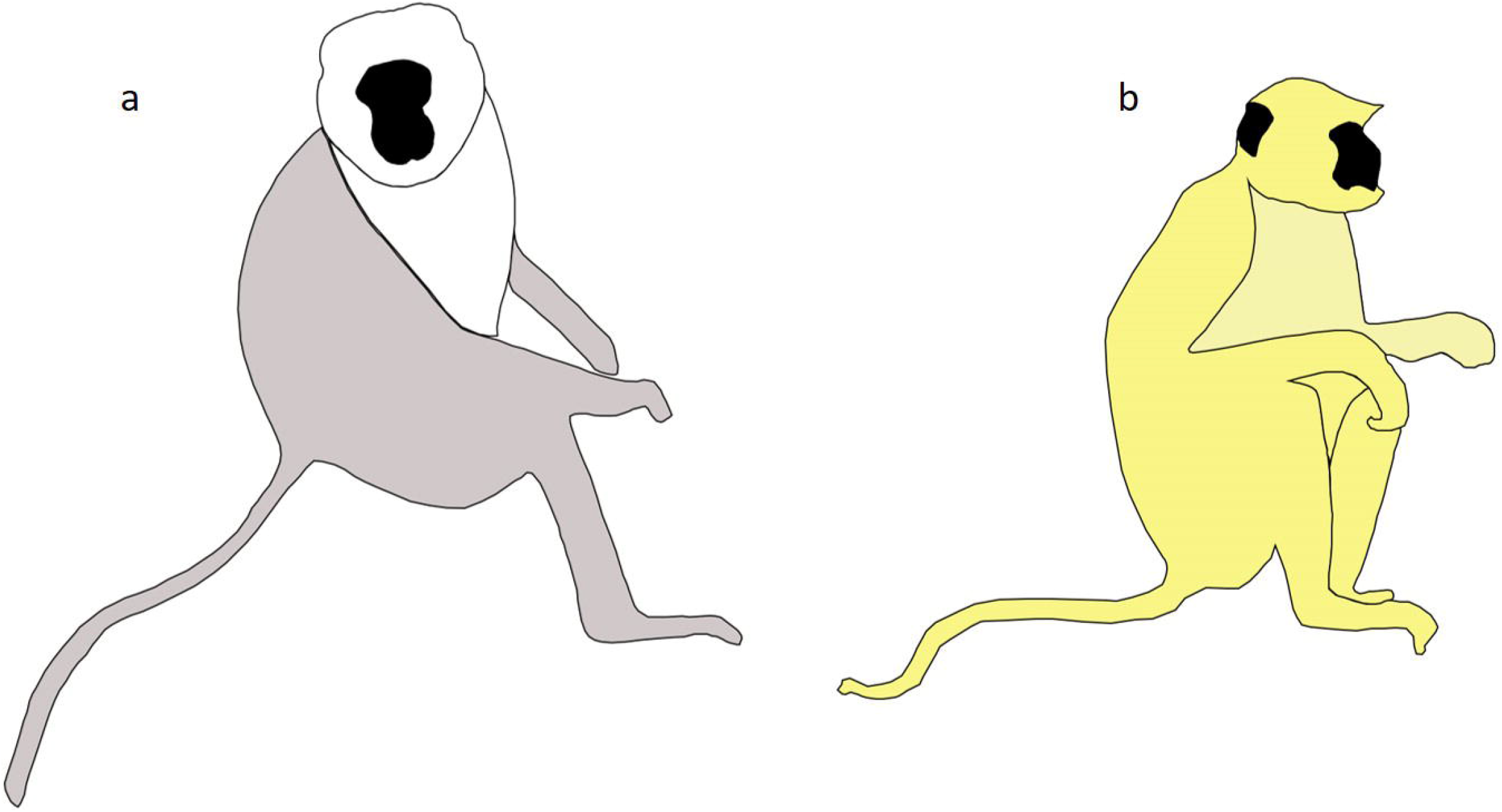
Contrast between the head and the dorsal region of the body in the Himalayan langur (a) and *Semnopithecus entellus* (b).

**Fig. 3:**
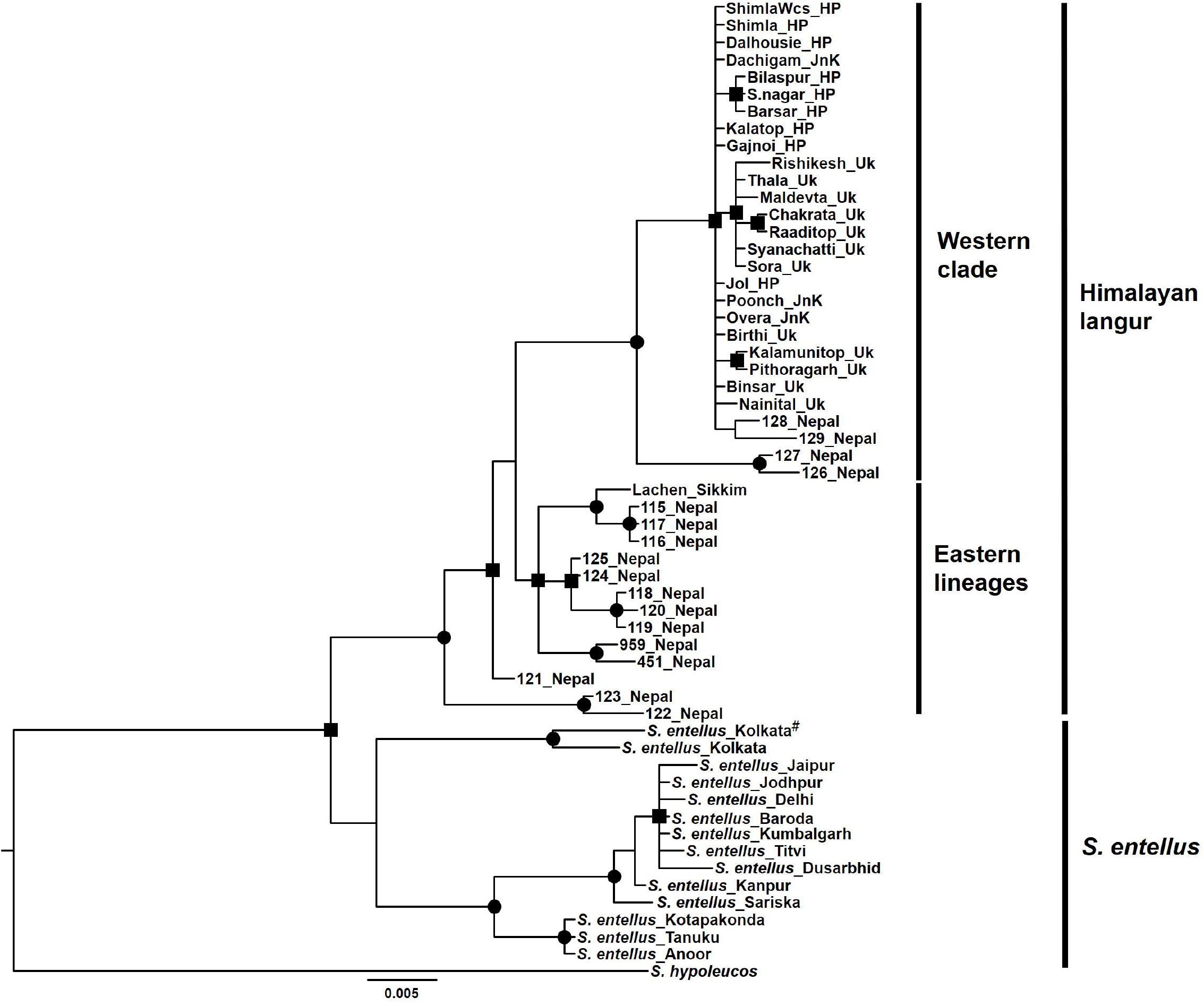
Bayesian phylogeny of Himalayan langur for mitochondrial cytochrome *b* (Cyt-b) gene. Node support is indicated by black circles and black squares. Black circle indicates high Bayesian posterior probability support (> 0.75) as well as a high maximum likelihood bootstrap support (>75). Black square indicates only a high Bayesian posterior probability support (> 0.75). HP = Himachal Pradesh; JnK = Jammu & Kashmir; Uk = Uttarakhand. ^#^Sequenced in this study.

### Hypothesis testing

The likelihood score of the best tree was significantly higher than that of the constraint tree for both SH test and AU tests. Therefore, these trees based on the molecular data did not support the splitting of Himalayan langurs into three species (Table 3).

**Table 3:** Topology test results.

| Tree | -ln L | Diff -ln L | SH | AU | p-value |
| --- | --- | --- | --- | --- | --- |
| Best tree | 2107.65046 | (best) |  |  |  |
| Constrained tree | 2371.94530 | 264.29484 | 0.0000* | ~0* | *P < 0.05 |

### Morphological data

Our final NJ tree (Fig. 4) was based on morphological data collected from 64 adult individuals from 60 localities. The NJ dendrogram retrieved a distinct cluster consisting of Himalayan samples that was sister to *S. entellus*. The UPGMA method also generated a similar topology.

**Fig. 4:**
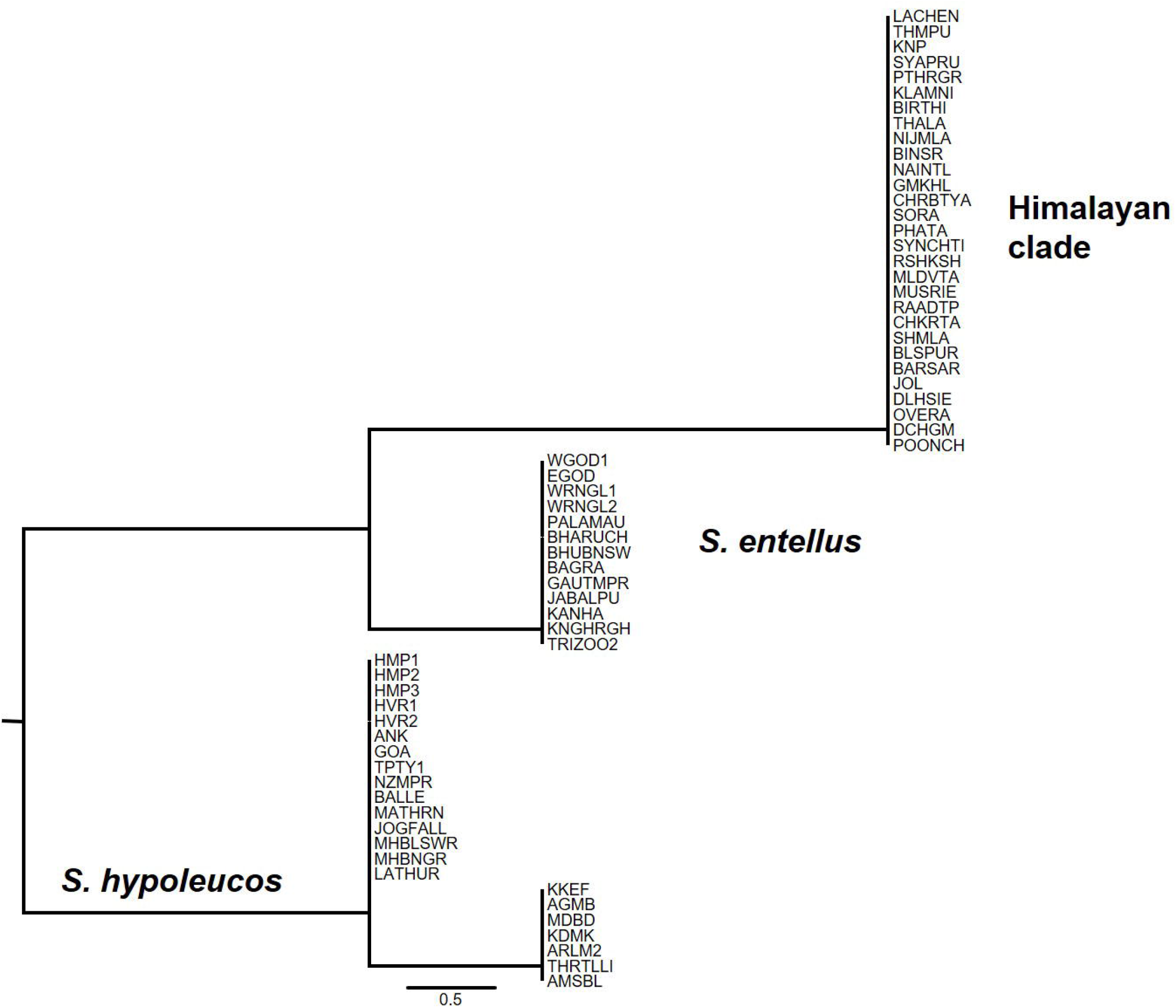
Neighbour-Joining tree based on 6 colour independent characters. *S. hypoleucos* is one of the two species of Hanuman langur from peninsular India

### Ecological niche modelling

#### Model prediction

The best model selected for the Himalayan langurs had features ‘LQPTH’ and a RM value of 2.5; whereas for *S. entellus*, the best model had features ‘Auto’ with a RM value of 3 (Table S4). Based on these models and the environmental variables selected, we obtained different distribution maps for *S. entellus* and the Himalayan langurs (Fig 5). In the distribution maps, the warmer colour indicates suitable area whereas the cooler colours indicate unsuitable areas. The AUC values for the training and test data for the Himalayan langur dataset were 0.9663 and 0.9621, respectively. And the AUC values for the training and test data for *S. entellus* were 0.889 and 0.87, respectively. These AUC values indicate that the potential distribution of these species fits well with our data.

**Fig. 5:**
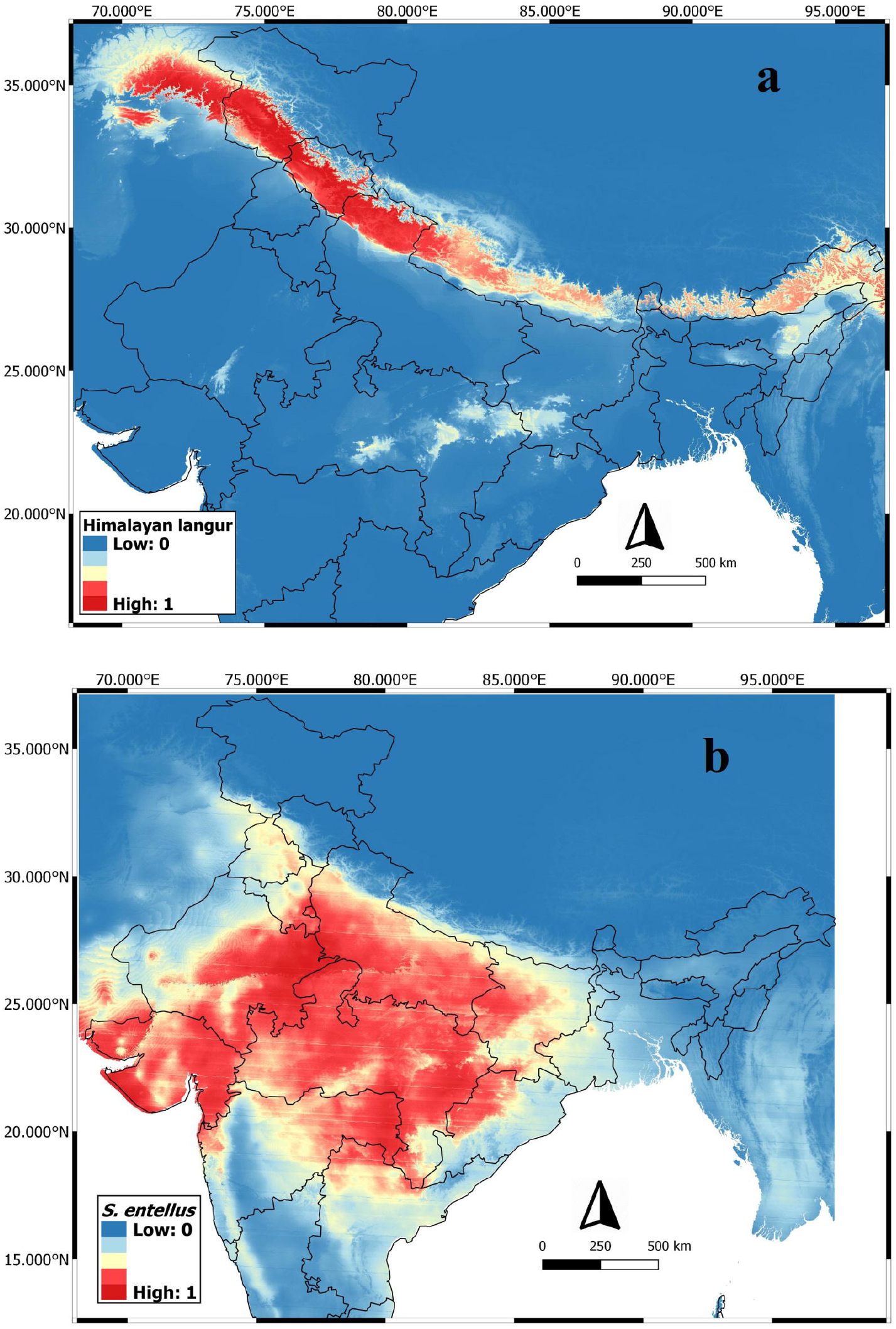
Species distribution maps of Himalayan langur (a) and *S. entellus* (b).

#### Variable selection and their importance

For the Himalayan langur, precipitation of driest quarter (Bio17) was the highest ranked variable (Table 5). Jackknife test also illustrates the importance of Bio17 (Supplementary material, Fig. S4). The response curve (Supplementary material, Fig. S2) shows that the habitat suitability increases with the precipitation of the driest quarter (Bio17) but eventually attains stationarity. Annual mean temperature (Bio1) and mean diurnal range (Bio2) were the next two contributing variables (Table 5).

Precipitation seasonality (Bio15) was the most contributing variable towards the distribution of *S. entellus* (Table 5). The response curve for Bio15 (Supplementary material, Fig S3) shows that the values for habitat suitability were high towards higher values of precipitation seasonality (Bio15), suggesting that *S. entellus* prefers areas with high variation in rainfall. Mean temperature of warmest quarter (Bio10) and annual mean temperature (Bio1) were the next two highly contributing variables (Table 5).

Niche overlap between *S. entellus* and Himalayan langur was 17 percent (Schoener’s D value = 0.17). However, when we compare the areas with high probability of distribution (> 0.75), there was no overlap.

## Discussion

The so-called Hanuman langur has been known to be a species complex since a long time. Nevertheless, recent studies have brought some clarity to their confused taxonomy. These studies suggest that the so-called Hanuman langur consists of at least three species: *S. entellus*, distributed in the plains of North India (Karanth et al., 2010); *S. priam* and *S. hypoleucos* distributed in peninsular India (Nag et al., 2011, Ashalakshmi et al., 2014, Nag et al., 2014). However, the taxonomic status of the Himalayan population of this complex remained unresolved. Here we address this issue by applying multiple lines of evidence to resolve the taxonomic status of the Himalayan langur.

Our molecular analysis based on mitochondrial Cyt-*b* gene retrieved a monophyletic Himalayan langur distinct from *S. entellus* of the plains. The Himalayan langur is also morphologically distinct and occupies a specific niche in the Himalayas. Thus, three lines of evidences, corresponding to different species concepts, suggest that the Himalayan langurs is separately evolving metapopulation lineage (de Queiroz, 1998). Phylogenetic species concept (PSC) II (de Queiroz, 1998; Donoghue, 1985; Mishler, 1985) identifies species as monophyletic groups on the basis of shared derived characters. Our genetic data shows that Himalayan langur and *S. entellus* are reciprocally monophyletic. Results from the morphological analysis suggests that the Himalayan langur is a separate species as per the Phenetic Species Concept (Sokal & Crovello, 1970). Our niche modelling analysis too shows that these two lineages occupy distinct ecological regions and thus show separation along the ecological axis (Van Valen, 1976).

Among the plethora of classification schemes only Hill (1939) placed the Himalayan langurs in a separate species, *S. schistaceus*, with multiple subspecies – *ajax, achilles, lanius* and *hector* (Table 1). Our study supports this classification, however there is no support for further splitting this species into multiple subspecies/species (see result under hypotheses testing). Therefore, we recommend *Semnopithecus schistaceus* Hodgson, 1840 as a species name for all the Himalayan langurs and subsume all the earlier described species and subspecies within it until further detailed taxonomic studies address this issue.

Two of the characters used in this study – HBC and forms of tail carriage (TC3 and TC4), can be used as field identification characters for differentiating *S. schistaceus* and *S. entellus*. We do not recommend the use any of the external morphological characters listed in the earlier classification schemes (Table 1) for distinguishing between different subspecies of Himalayan langurs. The morphological characters used in earlier classification schemes to describe these subspecies, are highly plastic, variable, and subjective in nature (Nag et al., 2011). For instance, in the identification key Pocock (1939) describes *schistaceus* as follows “General colour paler, salty or greyish-buff; coat shorter and less woolly” and for *achilles* he writes, “General colour dark earthy brown; coat thick and woolly”. Hill (1939) describes *schistaceus* as “A slatey-grey race, with shorter, less woolly coat than those found at higher altitudes”. These characters tend to differ based on what month of the year the langur is being observed and what is the altitude at that location. Roonwal (1981) describes four types of tail carriage in the NT langurs, however we recorded only two tail carriage types that are used in this study, the other two tail carriage types have not been observed.

We used ecological niche modelling (ENM) to determine the distribution range of *S. schistaceus*. It predicted the distribution range of the *S. schistaceus* (Fig. 5a) with high accuracy (AUC = 0.96). The distribution was mainly governed by precipitation of the driest quarter (Bio17). Response curves (Supplementary material, Fig. S2) for the top three contributing variables (Table 4) suggests that *S. schistaceus* prefers areas with high precipitation but with moderate temperatures. Interestingly, slope and aspect did not contribute to the model prediction. A recent study (Khanal et al., 2018) also showed that precipitation plays a major role in the distribution of *S. schistaceus* in Nepal. Langurs in the Himalaya inhabit broad leaves subtropical forest at lower altitudes and temperate broadleaf forest at higher altitudes (Bishop, 1979; Curtin, 1982; Minhas et al., 2013; Sayers & Norconk, 2008; Sugiyama, 1976). Occasionally, these langurs are found in temperate coniferous forests above 4000m altitude (Bishop, 1977). These forests receive high rainfall during the monsoon as well as precipitation in the form of snowfall in winter (Bhattarai & Vetaas, 2003; J. S. Singh & Singh, 1987; P. Singh, Ramasastri, & Kumar, 1995).

**Table 4:**
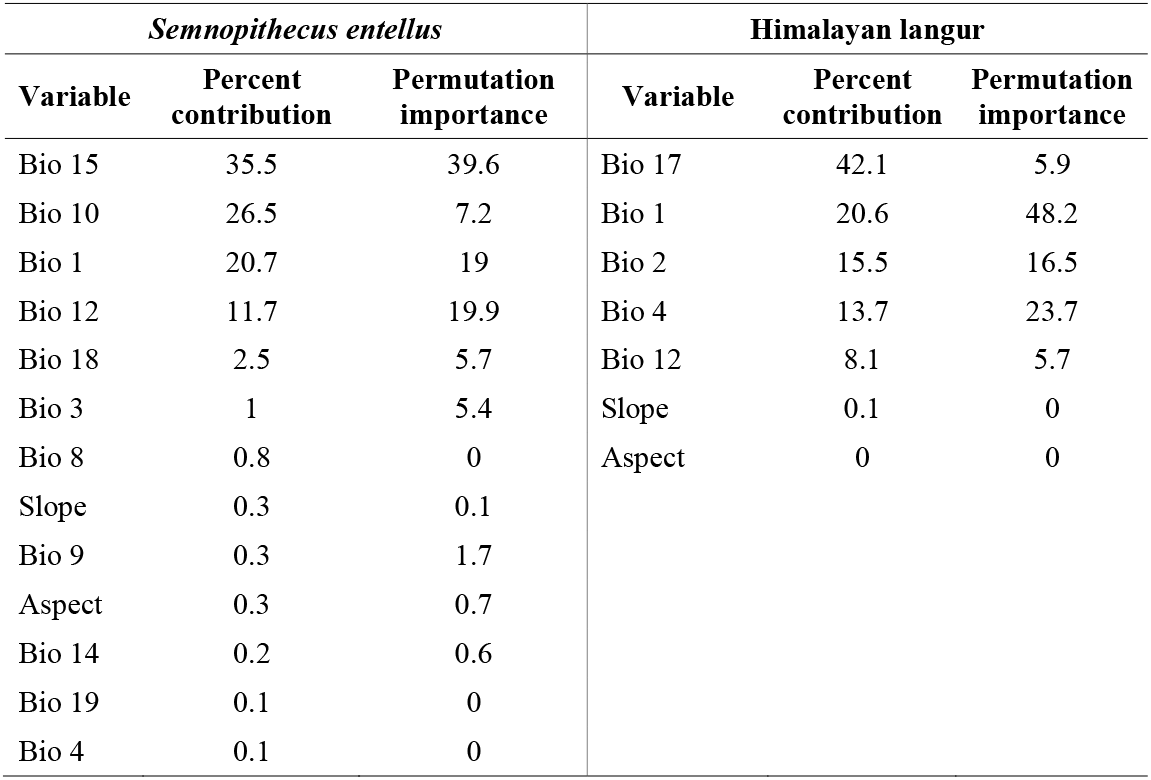
Maxent results showing the most important variables ranked on the basis of the amount of variation they explain in predicted distribution of *Semnopithecus entellus* and the Himalayan langur along with the permutation importance for the variables.

We also used ENM to determine if the ecological niches of *S. schistaceus* and *S. entellus* are separate. The ENM distinctly demarcated ecological niches of these two species. The AUC value for both the distribution models was significant, implying that the results greatly differ from the random predictions. Precipitation was the most important variable in demarcating these two species, with *S. entellus* requiring less precipitation and the Himalayan langurs need higher precipitation. Bishop (1979) also pointed out that climate is the factor that governs the distribution of the *S. schistaceus* and *S. entellus*. We also checked for niche overlap between these two species to determine if they are divergent in their ecological axis. The niche overlap between these two taxa was not significant suggesting that their ecological niches are separate.

This study provides a comprehensive evidence for elevating the Himalayan langur to a species that is distinct from *S. entellus* of the northern plains. Based on our results, we recommend that this species be assigned to *S. schistaceus* as per Hill (1939).

## Acknowledgments

We would like to thank the Department of Biotechnology, Government of India for providing funds for laboratory work. We would like to thank Dr. Chetan Nag and Ms. Mehreen Khaleel for providing some of the samples used in this study. Ms. Himani Nautiyal provided photographs from Bhutan for the morphological analysis. Dr. Bilal Habib for help during sample collection and Ms. Neha Tiwari for help in lab work. We would also like to thank the forest departments of Uttarakhand, Himachal Pradesh and Sikkim for providing necessary collection permits. Additionally, thanks to Jammu and Kashmir forest department for permits provided to Mehreen Khaleel. KA would like to thank the Rufford foundation and Primate Conservation Inc. (PCI) for providing funding for field work. Dr. Aravind Madhyastha and Mr. Bipin Charles for guidance in the niche modelling analysis.

