## Supplementary material for "Integrative taxonomy confirms the species status of the Himalayan langurs, *Semnopithecus schistaceus* Hodgson 1840"

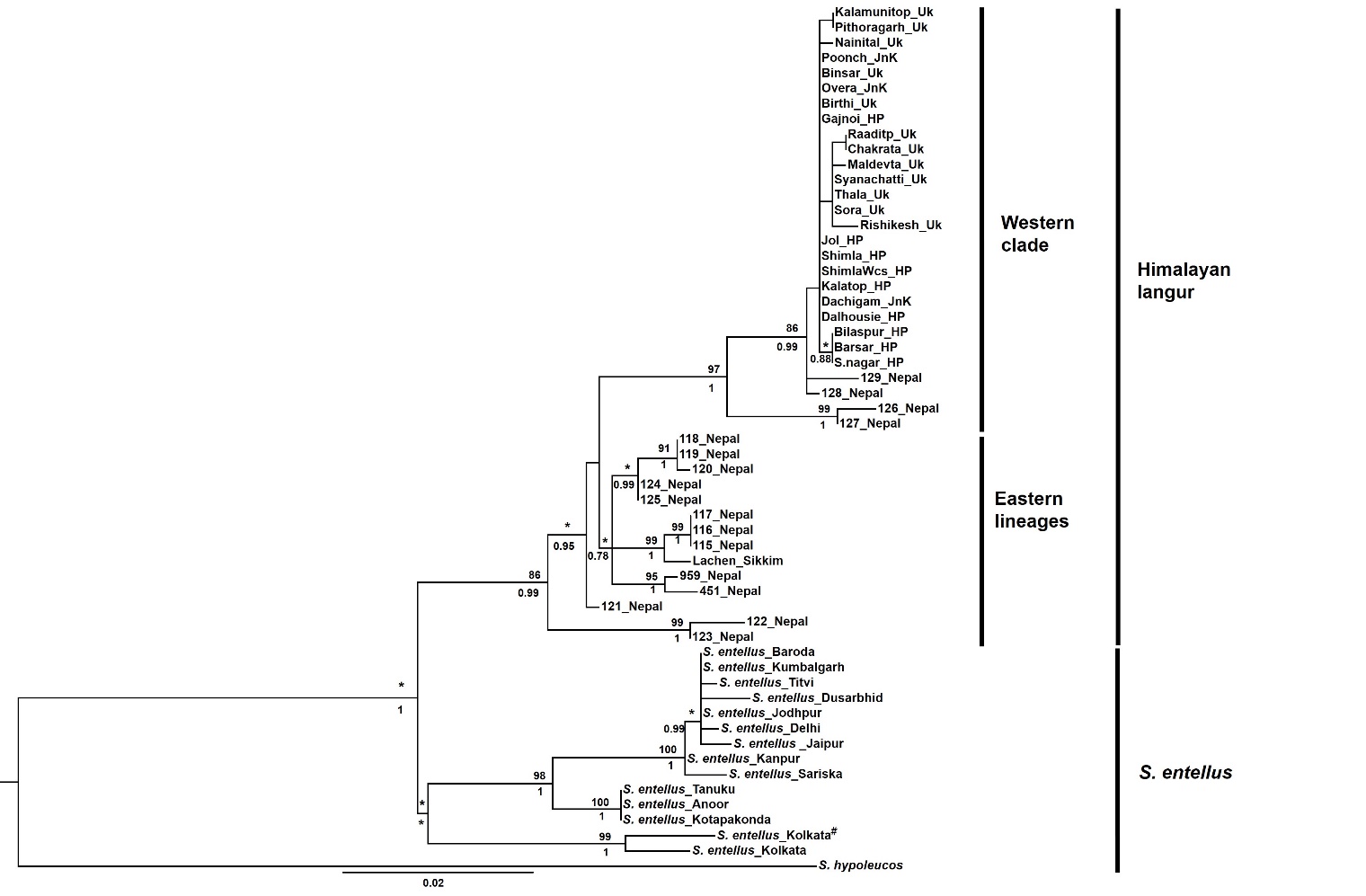

**Fig. S1:** Maximum likelihood tree of the Himalayan langur reconstructed using mitochondrial cytochrome – *b* (Cyt – *b*) gene. Number above and below the nodes are the bootstrap support values and posterior probability values, respectively, for each node. * indicates node support value < 75 or < 0.75. ^#^Sequenced in this study.

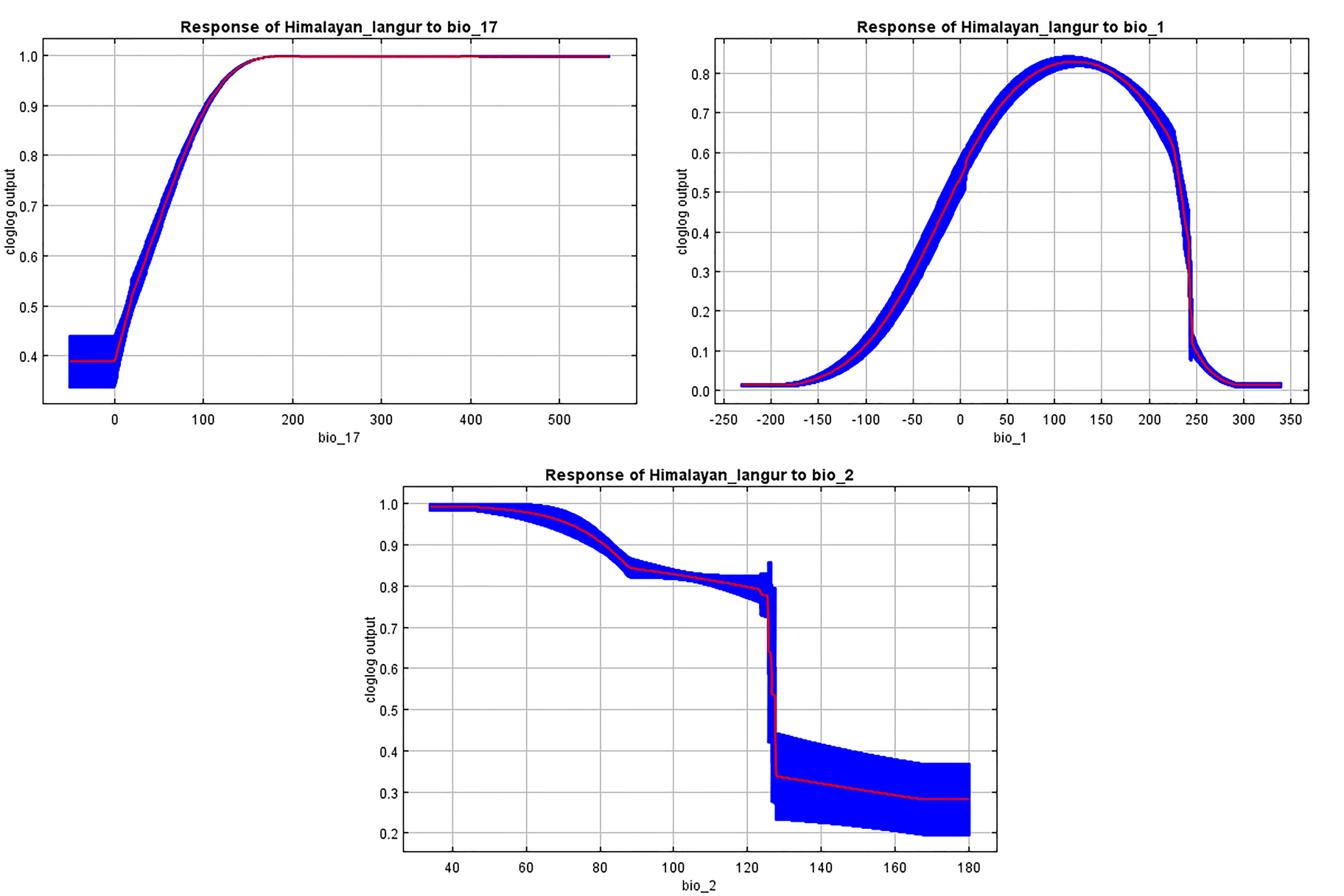

**Fig. S2:** Response curves for top three variables of importance in the Himalayan langur

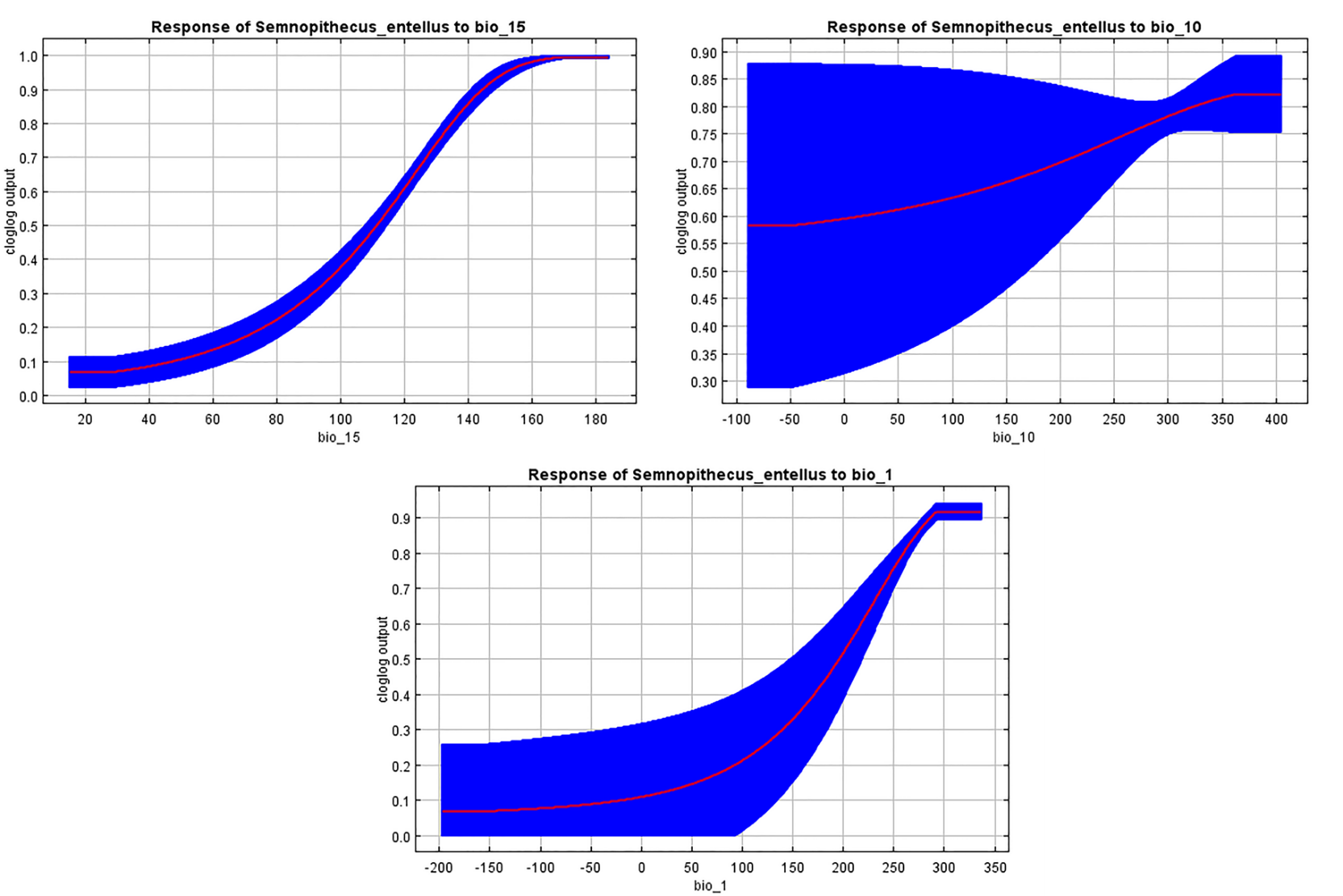

**Fig. S3:** Response curves for top three variables of importance in *Semnopithecus entellus*

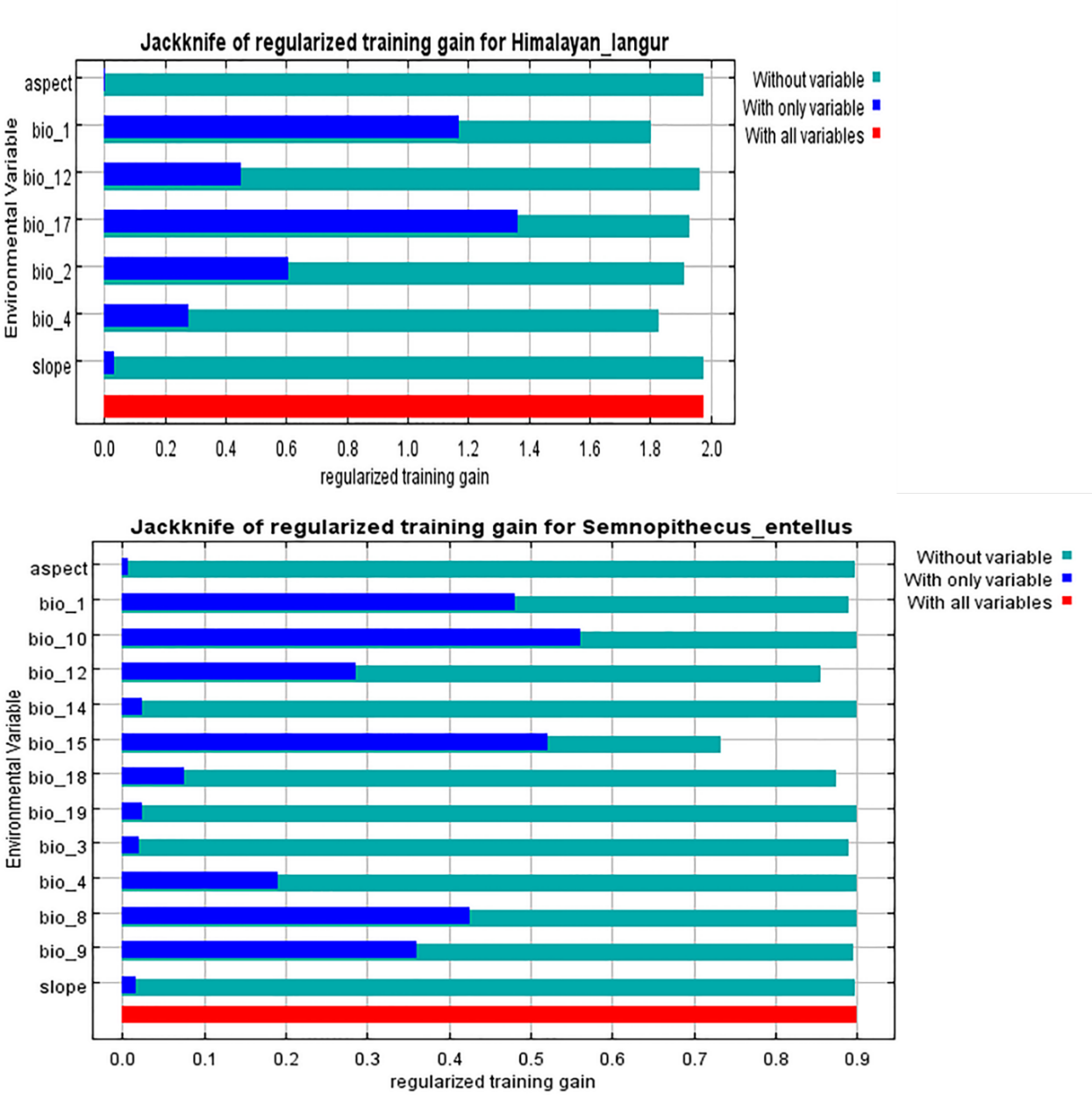

**Fig S4:** Jackknife test of regularised gain

**Table S1:** Locations of all the fecal samples collected for the study.

| **Latitude** | **Longitude** | **Elevation** | **No. of samples collected** |
| --- | --- | --- | --- |
| 33.68274 | 74.44421 | 3068 | 3 |
| 34.07608 | 74.56408 | 1796 | 6 |
| 34.07115 | 74.59235 | 2010 | 5 |
| 34.12110 | 74.99444 | 2087 | 7 |
| 33.95963 | 75.29607 | 2200 | 4 |
| 32.53807 | 75.95441 | 1817 | 1 |
| 32.53807 | 75.96306 | 1960 | 3 |
| 32.53407 | 75.97314 | 2031 | 4 |
| 32.53180 | 76.01758 | 2489 | 1 |
| 32.53511 | 76.04634 | 2269 | 6 |
| 32.52777 | 76.08361 | 1622 | 9 |
| 32.19456 | 76.13210 | 697 | 3 |
| 31.60419 | 76.43049 | 749 | 4 |
| 31.34576 | 76.78393 | 1025 | 3 |
| 31.34579 | 76.78499 | 993 | 1 |
| 31.50311 | 76.90276 | 1120 | 5 |
| 31.10310 | 77.15263 | 2091 | 3 |
| 31.10350 | 77.15423 | 2088 | 2 |
| 31.10409 | 77.15710 | 2106 | 5 |
| 31.10799 | 77.16646 | 2161 | 2 |
| 31.10026 | 77.23568 | 2321 | 1 |
| 30.68908 | 77.87001 | 2094 | 1 |
| 30.70000 | 77.87174 | 2041 | 5 |
| 30.46354 | 78.06320 | 2000 | 3 |
| 30.45103 | 78.08196 | 1831 | 2 |
| 30.33809 | 78.12877 | 764 | 5 |
| 30.77358 | 78.25728 | 2217 | 4 |
| 30.10681 | 78.29535 | 352 | 4 |
| 30.12418 | 78.31133 | 354 | 2 |
| 30.90474 | 78.36508 | 2012 | 5 |
| 30.05882 | 78.51107 | 593 | 1 |
| 30.76847 | 78.59856 | 1521 | 6 |
| 30.02100 | 78.63456 | 550 | 2 |
| 29.88449 | 78.67303 | 1550 | 3 |
| 30.38834 | 78.83901 | 2100 | 4 |
| 30.57578 | 79.04742 | 1587 | 6 |
| 29.31781 | 79.34721 | 619 | 1 |
| 29.34665 | 79.38558 | 1164 | 6 |
| 30.36582 | 79.46301 | 1476 | 4 |
| 30.05083 | 79.50986 | 1698 | 7 |
| 29.68749 | 79.73640 | 2004 | 6 |
| 30.03160 | 80.16652 | 1873 | 3 |
| 30.04284 | 80.19897 | 2609 | 5 |
| 29.66763 | 80.23032 | 1205 | 7 |
| 27.41350 | 88.19775 | 2129 | 1 |
| 27.75934 | 88.53851 | 2808 | 4 |

**Table S2:** Different environmental layers used in this study. Each layer is of 30arcsec resolution and is clipped to the region from 68 °E to 97.4 °E and from 6.7 °N to 37 °N.

| **Layer** | **Variable** | **Reference** | |
| --- | --- | --- | --- |
| Bio 1 | Annual Mean Temperature (°C*10) | [http://www.worldclim.org](http://www.worldclim.org/) | |
| Bio 2 | MeanDiurnalRange (Mean (period max-min)) (°C*10) | [http://www.worldclim.org](http://www.worldclim.org/) | |
| Bio 3 | Isothermality (Bio 2/Bio 7) (°C*10) | [http://www.worldclim.org](http://www.worldclim.org/) | |
| Bio 4 | Temperature Seasonality (SD*100) | [http://www.worldclim.org](http://www.worldclim.org/) | |
| Bio 5 | Max Temperature of Warmest month (°C*10) | [http://www.worldclim.org](http://www.worldclim.org/) | |
| Bio 6 | Min Temperature of Coldest month (°C*10) | [http://www.worldclim.org](http://www.worldclim.org/) | |
| Bio 7 | TemperatureAnnualRange (Bio 5-Bio 6) | [http://www.worldclim.org](http://www.worldclim.org/) | |
| Bio 8 | Mean Temperature of Wettest Quarter (°C*10) | [http://www.worldclim.org](http://www.worldclim.org/) | |
| Bio 9 | Mean Temperature of Driest Quarter (°C*10) | [http://www.worldclim.org](http://www.worldclim.org/) | |
| Bio 10 | Mean Temperature of Warmest Quarter (°C*10) | [http://www.worldclim.org](http://www.worldclim.org/) | |
| Bio 11 | Mean Temperature of Coldest Quarter (°C*10) | [http://www.worldclim.org](http://www.worldclim.org/) | |
| Bio 12 | Annual Precipitation (mm) | [http://www.worldclim.org](http://www.worldclim.org/) | |
| Bio 13 | Precipitation of Wettest Period (mm) | [http://www.worldclim.org](http://www.worldclim.org/) | |
| Bio 14 | Precipitation of Driest Period (mm) | [http://www.worldclim.org](http://www.worldclim.org/) | |
| Bio 15 | Precipitation Seasonality (Coefficient of Variation) | [http://www.worldclim.org](http://www.worldclim.org/) | |
| Bio 16 | Precipitation of Wettest Quarter (mm) | [http://www.worldclim.org](http://www.worldclim.org/) | |
| Bio 17 | Precipitation of Driest Quarter (mm) | [http://www.worldclim.org](http://www.worldclim.org/) | |
| Bio 18 | Precipitation of Warmest Quarter (mm) | [http://www.worldclim.org](http://www.worldclim.org/) | |
| Bio 19 | Precipitation of Coldest Quarter (mm) | [http://www.worldclim.org](http://www.worldclim.org/) | |
| DEM | Digital Elevation model | | USGS, EROS centre, Hydro1k for Asia |
| Aspect | Direction of slope | | USGS, EROS centre, Hydro1k for Asia |
| Slope | Difference between two neighbouring cells elevation | | USGS, EROS centre, Hydro1k for Asia |

**Table S3:** Correlation matrix between the 22 variables used in this study. Numbers below diagonal are for *S. entellus* and the numbers above diagonal shows values for the Himalayan langurs. Highlighted values indicate high correlation between two variables.

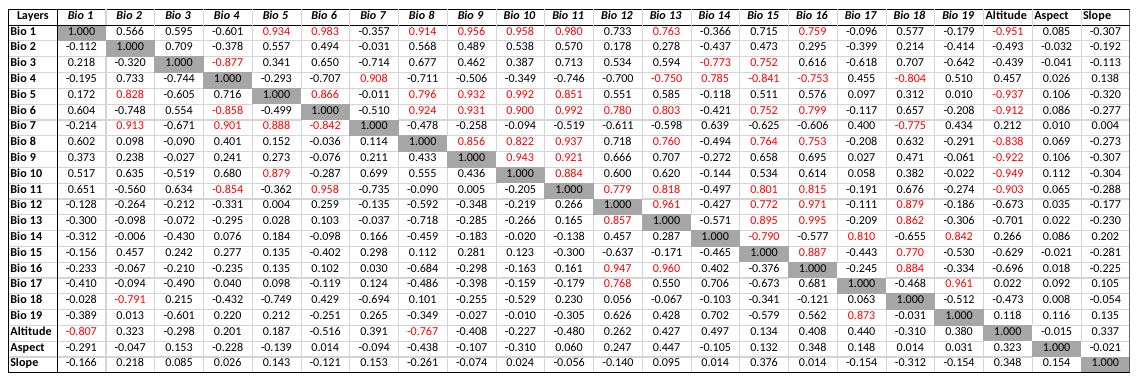

**Table S4:** Model selection for Maxent analysis: The table shows AUC values for different models. AUC values in bold shows the features and RM values selected for respective taxa.

| **Features** | **RM values** | | | | | | | | | | | |
| --- | --- | --- | --- | --- | --- | --- | --- | --- | --- | --- | --- | --- |
|  | **0.5** | **1** | **1.5** | **2** | **2.5** | **3** | **0.5** | **1** | **1.5** | **2** | **2.5** | **3** |
|  | **Himalayan langur** | | | | | | ***Semnopithecus entellus*** | | | | | |
| **Auto** | 0.962 | 0.958 | 0.968 | 0.966 | 0.963 | 0.963 | 0.861 | 0.869 | 0.877 | 0.859 | 0.867 | **0.88** |
| **L** | 0.893 | 0.872 | 0.877 | 0.867 | 0.878 | 0.888 | 0.837 | 0.844 | 0.838 | 0.849 | 0.856 | 0.846 |
| **LQ** | 0.957 | 0.958 | 0.955 | 0.955 | 0.958 | 0.958 | 0.846 | 0.862 | 0.86 | 0.846 | 0.858 | 0.859 |
| **LQP** | 0.961 | 0.961 | 0.963 | 0.965 | 0.961 | 0.965 | 0.87 | 0.842 | 0.848 | 0.847 | 0.846 | 0.851 |
| **LQPH** | 0.964 | 0.966 | 0.964 | 0.965 | 0.964 | 0.963 | 0.873 | 0.877 | 0.862 | 0.868 | 0.862 | 0.867 |
| **LQPTH** | 0.958 | 0.968 | 0.964 | 0.964 | **0.969** | 0.966 | 0.876 | 0.871 | 0.87 | 0.861 | 0.849 | 0.868 |
| **Q** | 0.862 | 0.851 | 0.853 | 0.855 | 0.871 | 0.863 | 0.844 | 0.847 | 0.849 | 0.846 | 0.837 | 0.847 |
| **T** | 0.961 | 0.96 | 0.961 | 0.963 | 0.958 | 0.958 | 0.867 | 0.868 | 0.857 | 0.845 | 0.832 | 0.819 |

L = Linear; Q = Quadratic, P = Product, H = Hinge; T = Threshold; RM = Regularisation Multiplier

**Table S5:** Sequences used for the phylogenetic analysis with their accession numbers, Sample IDs and location coordinates.

| **Sequences** | **Sample-ID** | **Latitude** | **Longitude** | **Accession Number** | **Sample type** |
| --- | --- | --- | --- | --- | --- |
| ShimlaWcs_HP | CES15308 | 31.10026 | 77.23568 |  | Fecal |
| Shimla_HP | CES13325 | 31.10409 | 77.1571 |  | Fecal |
| Dalhousie_HP | CES15324 | 32.53407 | 75.97314 |  | Fecal |
| Dachigam_JnK | CES13345 | 34.1211 | 74.9944 |  | Fecal |
| Bilaspur_HP | CES15311 | 31.34576 | 76.78393 |  | Fecal |
| S.nagar_HP | CES15313 | 31.50311 | 76.90276 |  | Fecal |
| Barsar_HP | CES15320 | 31.60419 | 76.43049 |  | Fecal |
| Kalatop_HP | CES15327 | 32.5318 | 76.01758 |  | Fecal |
| Gajnoi_HP | CES12317d | 32.5277 | 76.0836 |  | Fecal |
| *S*. *entellus*_Kolkata^#^ | CES15329 | 22.8822 | 88.3997 |  | Fecal |
| Rishikesh_Uk | CES13324 | 30.10681 | 78.29535 |  | Fecal |
| Jol_HP | CES13335 | 32.19456 | 76.1321 |  | Fecal |
| Poonch_JnK | CES18302 | 33.68274 | 74.44421 |  | Fecal |
| Overa_JnK | CES18304 | 33.95963 | 75.29607 |  | Fecal |
| Thala_Uk | CES17340 | 30.05083 | 79.50986 |  | Fecal |
| Birthi_Uk | CES17346 | 30.0316 | 80.16652 |  | Fecal |
| Kalamunitop_Uk | CES17351 | 30.04284 | 80.19897 |  | Fecal |
| Pithoragarh_Uk | CES17358 | 29.66763 | 80.23032 |  | Fecal |
| Binsar_Uk | CES17362 | 29.68749 | 79.7364 |  | Fecal |
| Maldevta_Uk | CES17384 | 30.33809 | 78.12877 |  | Fecal |
| Nainital_Uk | CES17366 | 29.34665 | 79.38558 |  | Fecal |
| Chakrata_Uk | CES17305 | 30.699997 | 77.87174 |  | Fecal |
| Syanachatti_Uk | CES17312 | 30.90474 | 78.36508 |  | Fecal |
| Raaditop_Uk | CES17316 | 30.77358 | 78.25728 |  | Fecal |
| Sora_Uk | CES17317 | 30.76847 | 78.59856 |  | Fecal |
| Lachen_Sikkim | CES18329 | 27.75934 | 88.53851 |  | Fecal |
| *S. hypoleucos* | CES09401 | 13.5074 | 75.0368 |  | Tissue |
| 115_Nepal |  |  |  | MH271115 |  |
| 117_Nepal |  |  |  | MH271117 |  |
| 116_Nepal |  |  |  | MH271116 |  |
| 125_Nepal |  |  |  | MH271125 |  |
| 121_Nepal |  |  |  | MH271121 |  |
| 124_Nepal |  |  |  | MH271124 |  |
| 118_Nepal |  |  |  | MH271118 |  |
| 120_Nepal |  |  |  | MH271120 |  |
| 119_Nepal |  |  |  | MH271119 |  |
| 123_Nepal |  |  |  | MH271123 |  |
| 122_Nepal |  |  |  | MH271122 |  |
| 127_Nepal |  |  |  | MH271127 |  |
| 128_Nepal |  |  |  | MH271128 |  |
| 126_Nepal |  |  |  | MH271126 |  |
| 129_Nepal |  |  |  | MH271129 |  |
| 959_Nepal |  |  |  | AF293959 |  |
| 451_Nepal |  |  |  | AY519451 |  |
| *S. entellus*_Jaipur |  |  |  | AF293957 |  |
| *S. entellus*_Kolkata |  |  |  | AF293958 |  |
| *S. entellus* _kanpur |  |  |  | JQ734760 |  |
| *S. entellus* _Jodhpur |  |  |  | JQ734761 |  |
| *S. entellus*_Sariska |  |  |  | JQ734691 |  |
| *S. entellus*_Delhi |  |  |  | JQ734690 |  |
| *S. entellus*_Baroda |  |  |  | JQ734726 |  |
| *S. entellus*_Kumbalgarh |  |  |  | JQ734727 |  |
| *S. entellus*_Titvi |  |  |  | JQ734724 |  |
| *S. entellus*_Dusarbhid |  |  |  | JQ734725 |  |
| *S. entellus­*_Kotapakonda |  |  |  | JQ734733 |  |
| *S. entellus*_Tanuku |  |  |  | JQ734702 |  |
| *S. entellus*_Anoor |  |  |  | JQ734701 |  |

^#^Sequenced in this study; *S* = *Semnopithecus*.

HP = Himachal Pradesh; JnK = Jammu & Kashmir; Uk = Uttarakhand.

**Table S6:** Details of the locations and characters used for the morphological tree.

| **Location** | **Code** | **Longitude** | **Latitude** | **Morphotype** | **Morphological characters** | | | | | |
| --- | --- | --- | --- | --- | --- | --- | --- | --- | --- | --- |
|  |  |  |  |  | **Crest** | **Streak** | **EOB** | **TL** | **TC** | **HBC** |
| East Godavari | EGOD | **^†^** | **^†^** | SE | 0 | 0 | 3 | 1 | 3 | 0 |
| West Godavari | WGOD1 | **^†^** | **^†^** | SE | 0 | 0 | 3 | 1 | 3 | 0 |
| Warangal | WRNGL1 | **^†^** | **^†^** | SE | 0 | 0 | 3 | 1 | 3 | 0 |
| Warangal | WRNGL2 | **^†^** | **^†^** | SE | 0 | 0 | 3 | 1 | 3 | 0 |
| Palamau | PALAMAU | **^†^** | **^†^** | SE | 0 | 0 | 3 | 1 | 3 | 0 |
| Bharuch | BHARUCH | **^†^** | **^†^** | SE | 0 | 0 | 3 | 1 | 3 | 0 |
| Bhubaneshwar | BHUBNSW | **^†^** | **^†^** | SE | 0 | 0 | 3 | 1 | 3 | 0 |
| Bagra | BAGRA | **^†^** | **^†^** | SE | 0 | 0 | 3 | 1 | 3 | 0 |
| Gautampur | GAUTMPR | **^†^** | **^†^** | SE | 0 | 0 | 3 | 1 | 3 | 0 |
| Jabalpur | JABALPU | **^†^** | **^†^** | SE | 0 | 0 | 3 | 1 | 3 | 0 |
| Kanha | KANHA | **^†^** | **^†^** | SE | 0 | 0 | 3 | 1 | 3 | 0 |
| Kangerghati | KNGHRGH | **^†^** | **^†^** | SE | 0 | 0 | 3 | 1 | 3 | 0 |
| Trivandrum | TRIZOO2 | **^†^** | **^†^** | SE | 0 | 0 | 3 | 1 | 3 | 0 |
| Hampi | HMP1 | **^†^** | **^†^** | SH1 | 0 | 1 | 3 | 2 | 1 | 0 |
| Hampi | HMP2 | **^†^** | **^†^** | SH1 | 0 | 1 | 3 | 2 | 1 | 0 |
| Hampi | HMP3 | **^†^** | **^†^** | SH1 | 0 | 1 | 3 | 2 | 1 | 0 |
| Haveri | HVR1 | **^†^** | **^†^** | SH1 | 0 | 1 | 3 | 2 | 1 | 0 |
| Haveri | HVR2 | **^†^** | **^†^** | SH1 | 0 | 1 | 3 | 2 | 1 | 0 |
| Ankola | ANK | **^†^** | **^†^** | SH1 | 0 | 1 | 3 | 2 | 1 | 0 |
| Bondla | GOA | **^†^** | **^†^** | SH1 | 0 | 1 | 3 | 2 | 1 | 0 |
| Tolpetty | TPTY1 | **^†^** | **^†^** | SH1 | 0 | 1 | 3 | 2 | 1 | 0 |
| Nizampur | NZMPR | **^†^** | **^†^** | SH1 | 0 | 1 | 3 | 2 | 1 | 0 |
| Balle | BALLE | **^†^** | **^†^** | SH1 | 0 | 1 | 3 | 2 | 1 | 0 |
| Matheran | MATHRN | **^†^** | **^†^** | SH1 | 0 | 1 | 3 | 2 | 1 | 0 |
| Jog, Gersoppa | JOGFALL | **^†^** | **^†^** | SH1 | 0 | 1 | 3 | 2 | 1 | 0 |
| Mahabhaleshwar | MHBLSWR | **^†^** | **^†^** | SH1 | 0 | 1 | 3 | 2 | 1 | 0 |
| Mahbubnagar | MHBNGR | **^†^** | **^†^** | SH1 | 0 | 1 | 3 | 2 | 1 | 0 |
| Lathur | LATHUR | **^†^** | **^†^** | SH1 | 0 | 1 | 3 | 2 | 1 | 0 |
| Kukke Subrahmanya | KKEF | **^†^** | **^†^** | SH2 | 0 | 1 | 4 | 2 | 1 | 0 |
| Agumbe | AGMB | **^†^** | **^†^** | SH2 | 0 | 1 | 4 | 2 | 1 | 0 |
| Moodbidre | MDBD | **^†^** | **^†^** | SH2 | 0 | 1 | 4 | 2 | 1 | 0 |
| Kudremukha | KDMK | **^†^** | **^†^** | SH2 | 0 | 1 | 4 | 2 | 1 | 0 |
| Aralam | ARLM2 | **^†^** | **^†^** | SH2 | 0 | 1 | 4 | 2 | 1 | 0 |
| Theerthahalli | THRTLLI | **^†^** | **^†^** | SH2 | 0 | 1 | 4 | 2 | 1 | 0 |
| Amasebail | AMSBL | **^†^** | **^†^** | SH2 | 0 | 1 | 4 | 2 | 1 | 0 |
| Poonch | POONCH | 33.68274 | 74.44421 | HyL | 0 | 0 | 0 | 1 | 4 | 1 |
| Dachigam | DCHGM | 34.1211 | 74.9944 | HyL | 0 | 0 | 0 | 1 | 4 | 1 |
| Overa | OVERA | 33.95963 | 75.29607 | HyL | 0 | 0 | 0 | 1 | 4 | 1 |
| Dalhousie | DLHSIE | 32.53407 | 75.97314 | HyL | 0 | 0 | 0 | 1 | 4 | 1 |
| Jol, Shahpur | JOL | 32.19456 | 76.1321 | HyL | 0 | 0 | 0 | 1 | 4 | 1 |
| Barsar | BARSAR | 31.60419 | 76.43049 | HyL | 0 | 0 | 0 | 1 | 4 | 1 |
| Bilaspur | BLSPUR | 31.34576 | 76.78393 | HyL | 0 | 0 | 0 | 1 | 4 | 1 |
| Shimla | SHMLA | 31.10409 | 77.1571 | HyL | 0 | 0 | 0 | 1 | 4 | 1 |
| Chakrata | CHKRTA | 30.7 | 77.87174 | HyL | 0 | 0 | 0 | 1 | 4 | 1 |
| Raaditop | RAADTP | 30.77358 | 78.25728 | HyL | 0 | 0 | 0 | 1 | 4 | 1 |
| Mussoorie | MUSRIE | 30.45103 | 78.08196 | HyL | 0 | 0 | 0 | 1 | 4 | 1 |
| Maldevta | MLDVTA | 30.33809 | 78.12877 | HyL | 0 | 0 | 0 | 1 | 4 | 1 |
| Rishikesh | RSHKSH | 30.10681 | 78.29535 | HyL | 0 | 0 | 0 | 1 | 4 | 1 |
| Syanachatti | SYNCHTI | 30.90474 | 78.36508 | HyL | 0 | 0 | 0 | 1 | 4 | 1 |
| Phata | PHATA | 30.57578 | 79.04742 | HyL | 0 | 0 | 0 | 1 | 4 | 1 |
| Sora | SORA | 30.76847 | 78.59856 | HyL | 0 | 0 | 0 | 1 | 4 | 1 |
| Chirbatiya | CHRBTYA | 30.38834 | 78.83901 | HyL | 0 | 0 | 0 | 1 | 4 | 1 |
| Gumkhal | GMKHL | 29.88449 | 78.67303 | HyL | 0 | 0 | 0 | 1 | 4 | 1 |
| Nainital | NAINTL | 29.34665 | 79.38558 | HyL | 0 | 0 | 0 | 1 | 4 | 1 |
| Binsar | BINSR | 29.68749 | 79.7364 | HyL | 0 | 0 | 0 | 1 | 4 | 1 |
| Nijmula | NIJMLA | 30.36582 | 79.46301 | HyL | 0 | 0 | 0 | 1 | 4 | 1 |
| Thala | THALA | 30.05083 | 79.50986 | HyL | 0 | 0 | 0 | 1 | 4 | 1 |
| Birthi | BIRTHI | 30.0316 | 80.16652 | HyL | 0 | 0 | 0 | 1 | 4 | 1 |
| Kalamunitop | KLAMNI | 30.04284 | 80.19897 | HyL | 0 | 0 | 0 | 1 | 4 | 1 |
| Pithoragarh | PTHRGR | 29.66763 | 80.23032 | HyL | 0 | 0 | 0 | 1 | 4 | 1 |
| Syapru Besi, Nepal | SYAPRU | 28.1766 | 85.3461 | HyL | 0 | 0 | 0 | 1 | 4 | 1 |
| Sachen | KNP | 27.4135 | 88.19775 | HyL | 0 | 0 | 0 | 1 | 4 | 1 |
| Lachen | LACHEN | 27.75934 | 88.53851 | HyL | 0 | 0 | 0 | 1 | 4 | 1 |
| Bhutan | THMPU | 27.5986 | 89.6280 | HyL | 0 | 0 | 0 | 1 | 4 | 1 |

**†** See Nag et. al. (2011) for longitude and latitude.

Morphotypes: SE = *S*. *entellus*; SH1 & SH2 = *S*. *hypoleucos*; HyL = Himalayan langur.

EOB = Extent of Blackness; HBC = Head Body contrast;

Crest, Streak and HBC, 0 = absent, 1 = present; TL = Tail loop, 1 = Northern type, 2 = Southern type; EOB till finger tips = 1, till knuckles = 2, till wrist = 3, till elbow = 4; TC = tail carriage, TC3 = 3, TC4 = 4, TC1 = 1.
